# Spatiotemporal temperature control by holographic heating microscopy unveils cellular thermosensitive calcium signaling

**DOI:** 10.1101/2025.01.09.632068

**Authors:** Kotaro Oyama, Ayumi Ishii, Shuhei Matsumura, Tomoko Gowa Oyama, Mitsumasa Taguchi, Madoka Suzuki

**Author notes:** **Corresponding Authors** Kotaro Oyama Takasaki Institute for Advanced Quantum Science, National Institutes for Quantum Science and Technology 1233 Watanukimachi, Takasaki-shi, Gunma 370-1292, Japan Madoka Suzuki Institute for Protein Research, Osaka University 3-2 Yamadaoka, Suita, Osaka 565-0871, JapanTel.

## Abstract

Optical microheating technologies have revealed how biological systems sense heating and cooling at the microscopic scale. Sensing is based on thermosensitive biochemical reactions that frequently engage membrane proteins, Ca^2+^ channels, and pumps to convert sensing information as the Ca^2+^ signaling in cells. These findings highlight the feasibility of thermally manipulating intracellular Ca^2+^ signaling. However, how the thermosensitive Ca^2+^ signaling would behave, particularly in multicellular systems, remains elusive. In this study, to extend the ability of the spatiotemporal temperature control by optical microheating technologies, we propose holographic heating microscopy. Water-absorbable infrared (IR) laser light is modulated by a reflective liquid crystal on a silicon spatial light modulator (LCOS-SLM). A computer-generated hologram displayed on the LCOS-SLM modulates the spatial phase pattern of the IR laser light to generate predesigned temperature gradients at the microscope focal plane. The holographic heating microscopy visualizes how thermosensitive Ca^2+^ signaling is generated and propagated in MDCK cells, rat hippocampal neurons, and rat neonatal cardiomyocytes. Moreover, the optical control of the temporal temperature gradient reveals the cooling-rate dependency of Ca^2+^ signaling in HeLa cells. These findings demonstrate the extended ability of holographic heating microscopy in investigating cellular thermosensitivities and thermally manipulating cellular functions.

## Introduction

Ca^2+^ signaling, a form of inter- and intracellular communication, is involved in various cellular processes such as metabolism, muscle contraction, and neural excitation ^1^. The spatiotemporal distribution of cytoplasmic [Ca^2+^] ([Ca^2+^]_i_) in Ca^2+^ signaling is controlled by a group of biomolecules such as ion channels, exchangers, pumps, and Ca^2+^-binding proteins. In biochemical reactions in general ^2^, the activities of these biomolecules can be thermally modulated. The temperature-sensitive transient receptor potential channel is a prominent example, which is activated within a narrow range of the environmental temperature ^3^. The rate of Ca^2+^ uptake by sarco-/endoplasmic reticulum Ca^2+^ ATPase (SERCA) is increased by heating and vice versa ^4,5^. Therefore, a combination of the thermosensitivities of the biomolecules involved in Ca^2+^ signaling could lead to intracellular thermosensitive Ca^2+^ signaling. However, it is difficult to expect in which manner the thermosensitivity would emerge.

To examine the thermosensitivities of living systems at the subcellular spatial scale, microscopic optical heating has been frequently employed ^6–8^. This is mainly because optical microheating technologies have two major advantages for this purpose. First, a micron-sized point heat source can generate a local temperature gradient. The thermal response of biomolecules and cells can be examined to a set of various amplitudes of heat stimulations simultaneously within the field of view of the fluorescence microscope. Second, the local temperature gradient can be generated and decayed within a second or shorter because of quick heat diffusion to the environment surrounding the point heat source. Transient localized heating is usually quicker than global heating of the whole specimen. Therefore, the thermal damage to biomolecules and cells caused by long exposure to heat stress can be minimized or even avoided.

A water-absorbable infrared (IR) light (wavelength, 1,455–1,480 nm) has been a major option because it is compatible with optical microscopy, including fluorescence Ca^2+^ imaging, and the temperature of targeted cells can be selectively controlled with a single-cell resolution ^9–12^. In these studies, the IR light was focused into a single spot through an objective lens, creating a single temperature gradient in the field of view. Despite its successful applications, optical heating microscopy still has the potential for further development, enabling more precise spatiotemporal control of local temperature and potentially revealing unexplored cellular sensing abilities. To achieve this, the method requires additional capabilities, such as (i) dynamically switching between multiple localized heat sources within the field of view without moving the specimen, (ii) a wider range of options for spatial temperature distributions, and (iii) a wider range of options for temporal temperature distributions.

Herein, we propose a holographic heating microscopy method based on the reflective liquid crystal on silicon (LCOS)-spatial light modulator (SLM). SLMs, such as the digital mirror device (DMD) and LCOS-SLM, can control light distributions in space ^13^. SLMs have been employed in biological applications with an IR laser. Wang et al. used DMD for the optical stimulation of cells expressing light-activated receptors ^14^. Shirasaki et al. introduced a scanner mirror to change the position of heating in a cell sorting chip ^15^. Compared with these SLMs, LCOS-SLMs can modulate the phase pattern of light with higher efficiency ^16^. The computer-generated hologram (CGH) displayed on LCOS-SLMs allows controlling the distribution of light at the microscopic focal plane. These holographic technologies have been widely applied in optical microscopy systems, such as optical tweezers ^17^, multisite two-photon excitation ^18^, rapid volumetric imaging ^19^, and high-throughput transfection ^20^. In the present study, we demonstrate its new abilities in spatiotemporally controlling the microscopic temperature. Ca^2+^ signaling was manipulated optothermally to reveal cellular responses to temporal temperature gradients.

## Results & Discussion

### Holographic heating microscope for spatially controlling the microscopic temperature

An LCOS-SLM was employed for the spatial and temporal control of the local temperature under the fluorescence microscope. First, a bitmap image, or “target image,” was prepared, which represents the heat source(s) in the focal plane of the microscope. Then, a CGH image for the target image was created (Figure 1A). Thereafter, water-absorbable IR laser light (λ = 1,475 nm) was irradiated to the LCOS-SLM displaying the CGH image. The modulated light was focused by a planoconvex lens (lens 1 in Figure 1A) to project the target image on a coverslip that was placed in the conjugate image plane of the microscope. An aluminum block was set on the coverslip to mask the 0^th^ diffraction light at the center of the projected image. The light passing through the coverslip was finally focused by an objective lens to generate the target image at the focal plane. The heating period was controlled by a mechanical shutter.

**Figure 1.**
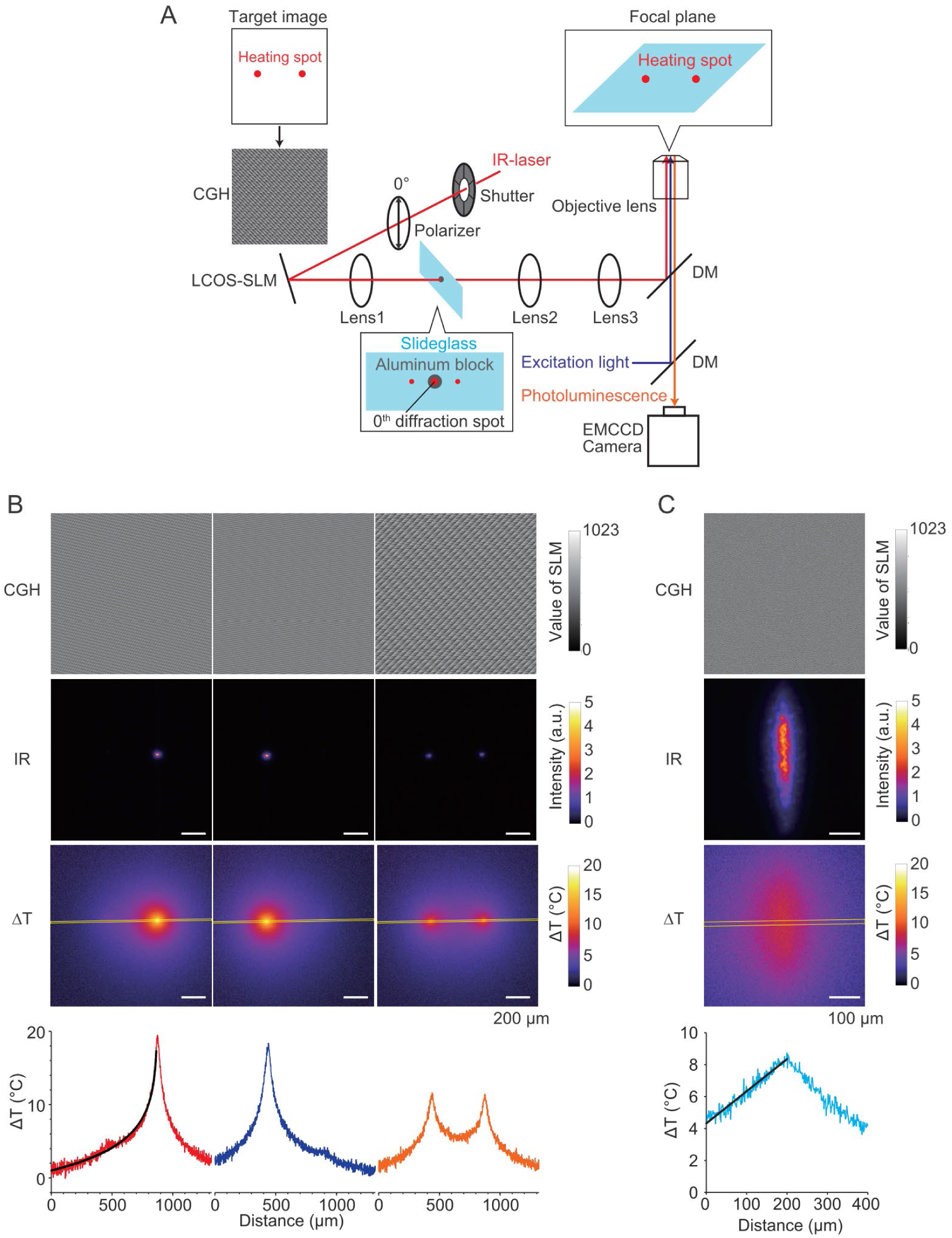
Spatial temperature control by the holographic heating microscope. (A) Optical setup for spatially controlling the temperature in the focal plane of the fluorescence microscope. A computer-generated hologram (CGH) image for a bitmap image with two dots was created. Water-absorbable infrared (IR) laser light (λ = 1,475 nm) was irradiated at a reflective liquid crystal on silicon (LCOS)-spatial light modulator (SLM) displaying the CGH images. The timing and duration of the irradiations were controlled by a mechanical shutter. The IR laser light polarization was aligned with the direction of the long axis (0 °) of the LCOS-SLM. The modulated light was focused on a coverslip by a planoconvex lens (lens 1) to generate the target image at the focal plane. The 0^th^ diffraction light that appeared as a bright spot at the center of the generated image was cut by an aluminum block on the coverslip. The light passing through the coverslip was adjusted by two planoconvex lenses (lenses 2 and 3) and focused by an objective lens to generate the target image at the focal plane of the microscope. For the imaging of the luminescent thermometer nanosheets and Ca^2+^ signaling, an LED source and an EMCCD camera were used. (**B**) Top images show CGH images for single-spot heating on the right or left side or double-spot heating. Middle images show the luminescence images of an IR-viewer sheet with upconversion nanoparticles on a coverslip. The bottom images show the temperature mapping by a luminescence thermometer nanosheet on a polymer-bottom dish. Graphs in the bottom show the temperature gradients analyzed. The temperature profiles along the yellow rectangles (width, 12.9 µm) were analyzed. The black curve was a fitting by Δ*T* = −3.67 ln(*d*/*d*_0_) + 25.9 (°C), where *d* is the distance from the heat source (µm), and *d*_0_ = 1 (in µm). *T*_0_, 30°C. Laser power, 12 mW. Scale bars, 200 µm. (**C**) Same as (**B**) but for line-shaped heating. The black line is a fitting by Δ*T* = −0.02 *d* + 8.35 (°C). *T*_0_, 30°C. Laser power, 8 mW. Scale bars, 200 µm.

As a simple demonstration of holographic heating, single-spot heating at two different locations, double-spot (Figure 1B) heating, and line heating were examined (Figure 1C). The target image generated by the IR laser light at the microscope focal plane was visually confirmed by an IR-viewer sheet with upconversion nanoparticles (ϕ = 100–300 nm, *λ*_ex_ = 1,550 nm, *λ*_em_ = 528, 542, and 662 nm) ^21^ (Figure S1, Movie S1). The actual temperature distributions visualized by the luminescent thermometer nanosheet appeared to be broader than the respective IR distributions because of thermal diffusion. By switching two CGH images of a single spot at different locations, temperature distributions were formed alternatively at different locations on the focal plane (Movie S2). For spot heating, the spatial gradient of the temperature increase, Δ*T*, was a linear function of ln(*d*) (Δ*T* = A − B ln(*d*)), where *d* is the distance from the heat source, and A and B are constants. Therefore, the shape of the heat source can be considered cylindrical rather than spherical, as we have previously reported ^22,23^. For line heating, Δ*T* was a linear function of *d* (Δ*T* = A − B*d*), as expected from the geometry. Thus, the holographic heating microscope can spatially control the temperature gradient at the focal plane.

### Spatial manipulation of Ca^2+^ signaling

Holographic heating microscopy was applied to examine cell type-dependent Ca^2+^ signaling in cell populations. We previously reported heat-induced Ca^2+^ bursts in the human fibroblast cell line WI-38 during optical heating (Δ*T* > ∼10°C) ^22^ and proposed the following mechanism: (i) heating decreases [Ca^2+^]_i_ and increases [Ca^2+^] in ER ([Ca^2+^]_ER_) quickly because of the acceleration of Ca^2+^ uptake by Ca^2+^ pump, SERCA ^4,5^ combined with the decrease in the open probability of Ca^2+^ channels, IP_3_R, as well as RyR ^24,25^. (ii) As the gradient of the electrochemical potential of Ca^2+^ between [Ca^2+^]_i_ and [Ca^2+^]_ER_ is enhanced, the Ca^2+^ uptake by SERCA gradually decreases, and the Ca^2+^ outflow from the ER to the cytoplasm through IP_3_R is increased. (iii) Finally, the Ca^2+^ outflow induced further Ca^2+^ release through IP_3_R because of the mechanism of Ca^2+^-induced Ca^2+^ release (CICR). The results of that study suggest that such a thermosensing system mediated by SERCA and IP_3_R would be widely observed in various mammalian cells.

To test this idea, we used Madin–Darby canine kidney (MDCK) cells as a representative epithelial cell line that is frequently employed in cell biology. MDCK cells were stimulated at two locations alternately with 10 s heating and 10 s blank by switching the CGH images of single-spot heating (Figure 1B). As expected, local heating induced Ca^2+^ bursts in the MDCK cells at the target position locally (Figure 2A and 2B, Movie S3). Then, second heating was applied at the other coordinates in the focal plane, inducing another local Ca^2+^ burst at the location where the second heat source was created. The third heating in the coordinates where the first heating was performed again induced Ca^2+^ burst in the similar cell population but with less amplitude than that in the first heating.

**Figure 2.**
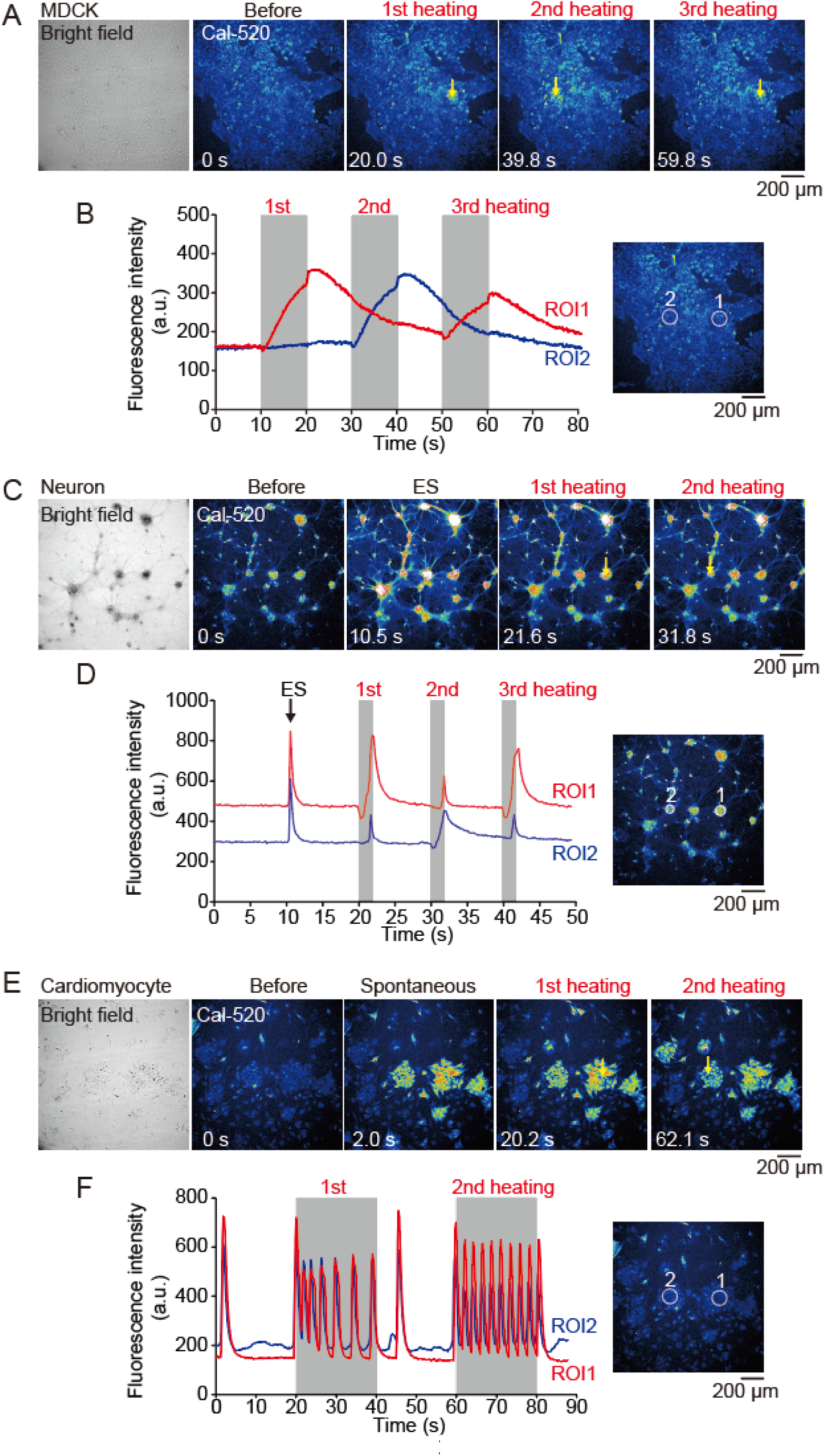
Optical manipulation of Ca^2+^ signaling by the holographic heating microscope. **(A)** Bright-field and fluorescence images of MDCK cells stained with the fluorescent Ca^2+^ indicator Cal-520. The first, second, and third heatings for 10 s were initiated at 10, 30, and 50 s, respectively, at the locations indicated by yellow arrows. *T*_0_, 37°C. (**B**) Left; time courses of the fluorescence intensities of Cal-520 in MDCK cells responding to optical heating for 10 s. Intensities in two regions of interest (ROIs 1 and 2) shown in the right image were analyzed. ROI1 was targeted in the first and third heatings, and ROI2 was targeted in the second heating. (**C**) Bright-field and fluorescence images of rat hippocampal neurons stained with Cal-520. Electrical stimulation (ES) was applied at 10 s. The first, second, and third heatings for 2 s were initiated at 20, 30, and 40 s, respectively, at the positions indicated by yellow arrows. *T*_0_, 37°C. (**D**) Left; time courses of the fluorescence intensities of Cal-520 in neurons responding to ES or optical heating for 2 s. ROI1 was targeted in the first and third heatings, and ROI2 was targeted in the second heating. The right images show ROI1 and ROI2. (**E**) Bright-field and fluorescence images of rat neonatal cardiomyocytes stained with Cal-520. The first and second heatings for 20 s were initiated at 20 and 60 s, respectively, at the positions indicated by the yellow arrows. *T*_0_, 26°C. (**F**) Left; time courses of the fluorescence intensities of Cal-520 in cardiomyocytes responding to optical heating for 20 s. ROI1 and ROI2 were targeted in the first and second heatings. The right images show ROI1 and ROI2. Scale bars, 200 µm. The gray bars in **B,D, and F** indicate the period of the maximum heating. Δ*T* at the center of the heating points was 18°C–19°C (Figure 1B).

The temperature changes affect the properties of the fluorescent Ca^2+^ indicators in general. In the current case, heating could decrease the Cal-520 intensity because of thermal quenching or increase its intensity of resting [Ca^2+^]_i_ (<100 nM) due to elevated Ca^2+^ affinity, as in the case of fluo-4 ^26^. However, the temporal changes in intensities were decoupled from those in temperature (Figure 2B). The gradual rise and return of the Cal-520 intensity were noted with a delay from the initiation and cessation of heating, respectively. Therefore, the increase in the Cal-520 intensity during heating represents the [Ca^2+^]_i_ increase.

Then, rat hippocampal neurons and neonatal cardiomyocytes were examined. Both cell types communicate via synchronized Ca^2+^ signaling through electrical coupling. Therefore, the thermosensitive Ca^2+^ signaling generated at the heat source is expected to propagate to other regions rapidly (quicker than the order of seconds), which is distinct from the case observed in MDCK cells. Heating of neurons for 2 s, which have been cultured for 21 days until a cellular network was formed, induced Ca^2+^ transients in a manner that is comparable to that induced by electrical stimulation (Figure 2C and 2D, Movie S4). Ca^2+^ transients were observed in nearly all cells in the field of view synchronously, indicating that the heat pulses excited the targeted neurons, and the excitation propagated through the cellular network. Either one or a combination of the following mechanisms can be considered: (i) activation of thermosensitive ion channels such as TRPV1 and TRPV2 ^3^, (ii) increase in membrane capacitance due to the thermal– mechanical effect ^27–29^, and (iii) activation of mechanosensitive ion channels ^30^ by convection flow during local heating ^23,31,32^. Further studies with electrical recording and antagonists for thermo- or mechanosensitive ion channels are required to reveal the details of the mechanism.

Lastly, the synchronized response was also observed in rat neonatal cardiomyocytes cultured for 4 days after isolation (Figure 2E and 2F, Movie S5). A group of cells at the heat source (ROI1) responded to the heating immediately, i.e., the frequency of intracellular Ca^2+^ transient was increased during heating as observed previously ^33^, and cells in the group responded simultaneously. Furthermore, the Ca^2+^ response was induced in other nearby groups. Changing the heating location to one of these groups (ROI2) induced a response similar to the primary group (ROI1). The synchronized behavior of these groups indicates the cellular tight coupling between these cells as in neurons.

In summary, the spatial manipulation of the local temperature was employed to demonstrate heat-induced Ca^2+^ signaling. Temperature is one of the major parameters governing the functions of the brain and heart. For example, the daily spatiotemporal rhythm in the human brain temperature is related to the age, health, and sex of the individual, and it affects the survival rate from brain injury ^34^. We previously reported that optical heating induced cardiac contraction without Ca^2+^ transients through the thermal activation of thin filaments ^10,35,36^. Therefore, the holographic heating microscope is potentially a useful tool to investigate the effects of temperature on the functions of the brain and heart in ex vivo and in vivo and further to manipulate them remotely.

### Holographic heating microscope for temporally controlling the microscopic temperature

As the second application of LCOS-SLM, an optical system was designed for temporally controlling the local temperature (Figure 3A). There has been no control of the cooling rate as the cooling process relies on heat diffusion. In the proposed system, two linear polarizers were placed at the paths before and after LCOS-SLM. The first polarizer was aligned in the 45° direction against the long axis (0°) of the LCOS-SLM. The second polarizer was set to cut off the IR light that was reflected without phase modulation by the LCOS-SLM. Here, CGH images were created to modulate the phase difference of the IR light. The modulated light passes through the second polarizer in accordance with the following equation: *I* = *I*_max_ (sinΘ)^2^ where *I* and *I* _max_ are the light intensity passed through and the maximum intensity, respectively, and Θ is the phase difference induced by the LCOS-SLM. When the values of the CGH images displayed on the LCOS-SLM were modulated sequentially from 416 to 0, IR light polarization was changed from linear (parallel to the second polarizer with π/2 of the phase difference; 100% transparency), elliptical, circular (π/4 of the phase difference; 50% transparency), elliptical, and at the end returned to linear (perpendicular to the second polarizer with no phase difference; 0% transparency). In the present study, 14 CGH images with a fixed gap value of 32 from full (value 416) to null (value 0) transmissions were created (Figure 3B). Then, the cooling rate was controllable by the dwell time between images (Figure 3B). A representative set of rapid, middle, and slow rates of cooling at the center of the heating spot were achieved, approximately corresponding to −5, −2, and −1 **°**C/s, respectively, (Figure 3C).

**Figure 3.**
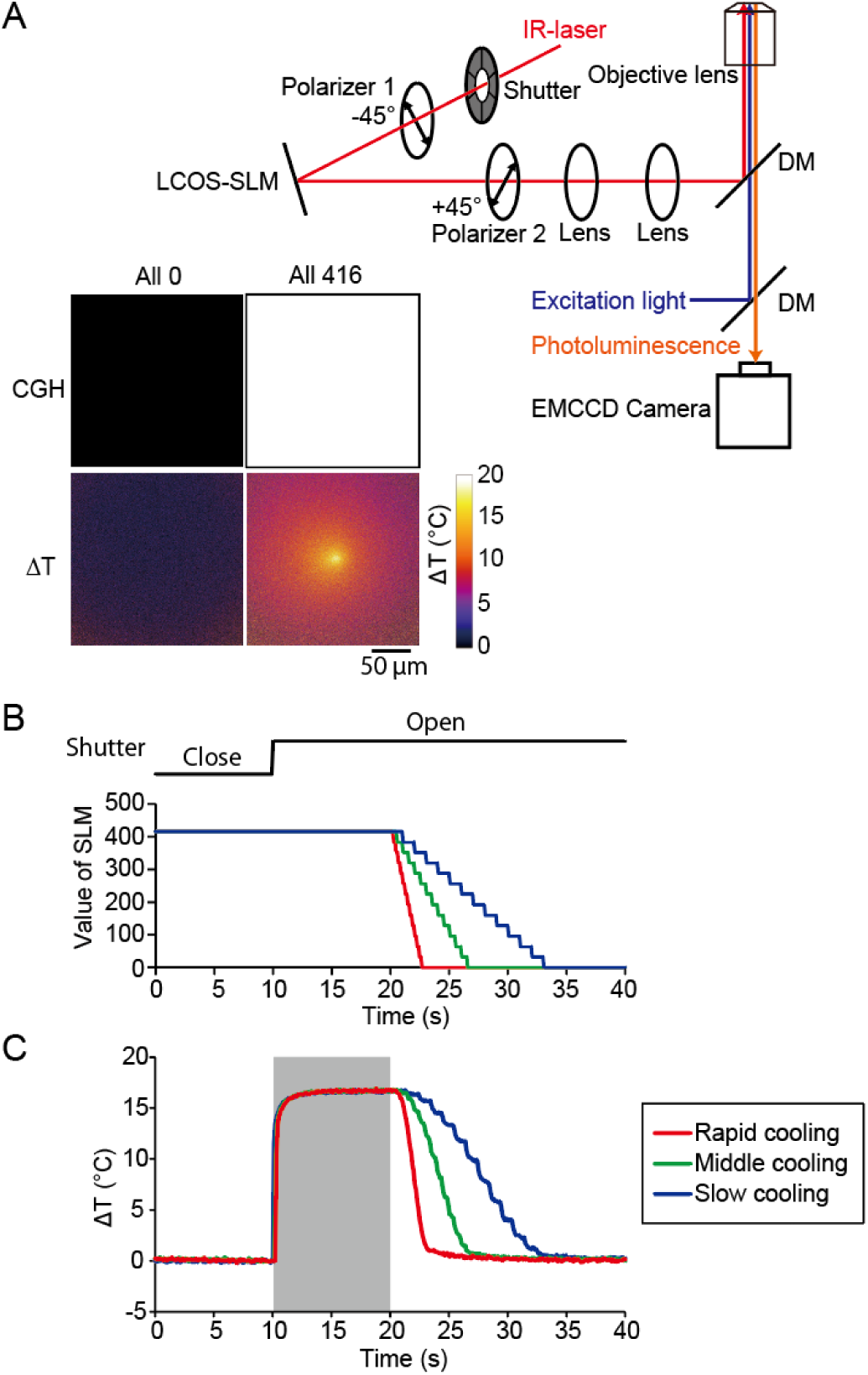
Temporal temperature control by the holographic heating microscope. (**A**) Optical setup for the temporal temperature control. Polarizer 1 was aligned in the 45° direction against the long axis of the LCOS-SLM. Polarizer 2 was set to cut off the IR light without modulation by LCOS-SLM. LCOS-SLM modulates the phase of light in the direction of the long axis of LCOS-SLM. The full-phase inversion gives the maximum transmission through polarizer 2. CGH images show the null (left) and full (right) transmissions of IR light and the corresponding temperature gradients at the microscopic focal plane measured with a luminescent thermometer nanosheet on a glass-bottom dish. The intensities at each pixel for null and full were 0 and 416, respectively. (**B**) Experimental processes for the optical manipulation of the temporal changes in the local temperature. The initial intensity of each pixel in the CGH image was set to 416. Opening a mechanical shutter initiates heating at the focal plane. Following exchanges of 13 CGH images cooled gradually. The periods of exchanges were 0.2, 0.5, and 1.0 s for rapid, middle, and slow cooling, respectively. (**C**) Temperature changes (Δ*T*) at the center of heating. The area within 5.5 µm from the center was analyzed. Gray bars indicate the period of the maximum heating. *T*_0_, 37°C. Laser power, 13 mW.

### Induction of the Ca^2+^ burst is governed by the cooling rate

Finally, temporal temperature modulation was applied to investigate the Ca^2+^ burst induced by rapid cooling. Rapid cooling in muscle cells induces Ca^2+^ release from the SR via ryanodine receptors, and muscle contraction is induced, which is called rapid cooling contracture (RCC) ^37^. The amplitude of the RCC is dependent on the cooling rate ^38^. Similar cooling-rate sensitivity in Ca^2+^ bursts was reported in plant cells ^39,40^. We previously reported that a transient heating (heat pulse) for 2 s and subsequent cooling induces Ca^2+^ bursts in HeLa cells ^31^. The proposed mechanism in HeLa cells was that sudden exposure to temperature causes an imbalance in Ca^2+^ homeostasis and Ca^2+^ release from the ER to the cytosol via the IP_3_R receptors. Inspired by these reports, in the present study, we examined the cooling rate dependency of the Ca^2+^ burst in HeLa cells. We heated the HeLa cells for 30 s and then exposed them to recooling with different cooling rates (Figure 4; Movies S6–S8). Transient Ca^2+^ bursts were observed in some cells during heating, and the subsequent rapid- or middle-rate cooling induced Ca^2+^ bursts immediately after the cessation of heating. However, no significant Ca^2+^ bursts were noted after the slow-rate cooling (Figure 4E). These results demonstrate that the Ca^2+^ bursts induced by heat pulses were dictated at the cooling rate.

**Figure 4.**
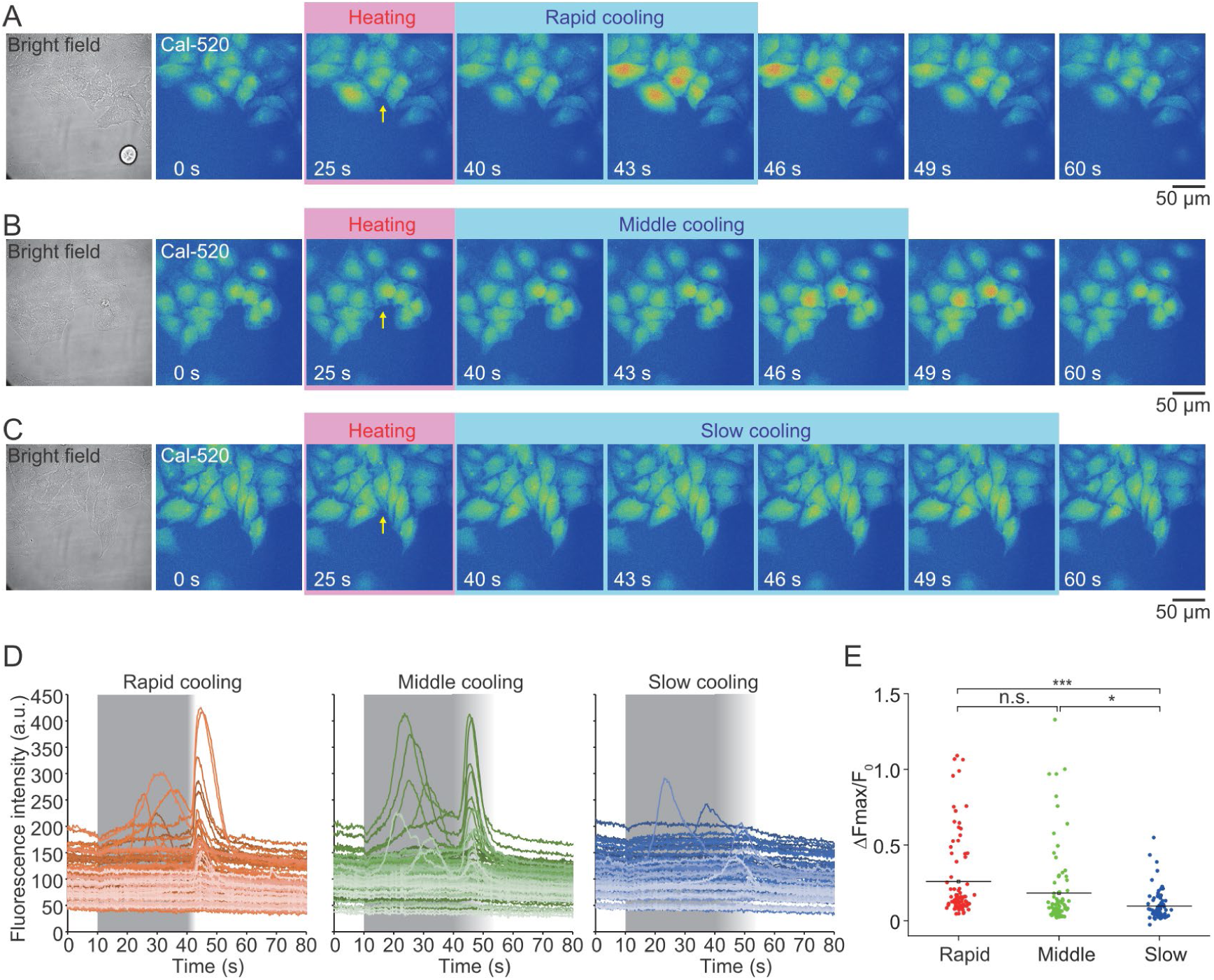
The induction of Ca^2+^ burst depends on the cooling rate. (**A–C**) Bright-field and fluorescence images of HeLa cells stained with the fluorescent indicator Cal-520. Optical heating for 30 s was initiated at 10 s. The yellow arrows indicate the locations where the IR light was focused. At 40 s, the cellss were exposed to the recooling protocol with rapid (**A**), middle (**B**), or slow (**C**) rates (Figure 3C). Scale bars, 50 µm. (**D**) Time courses of the fluorescence intensities of Cal-520 in HeLa cells responding to the thermal stimulation with rapid (left), middle (center), or slow (right) recooling rates. Gray bars indicate the period of the maximum heating. (**E**) The maximum changes in the intensities of Cal-520 induced by recooling. The maximum changes during recooling (Δ*F*_max_) were normalized to the intensities for 2 s at the end of heating (*F*_0_). The cell counts for rapid, middle, and slow recooling were 87, 88, and 77, respectively. The amplitudes were compared with the Tukey–Kramer test. * P < 0.05; *** P < 0.001; n.s., not significant (P > 0.05). *T*_0_, 37°C; Δ*T*, 8±1°C (mean±SD).

Based on current and previous results, the cooling rate dependency of the heat-induced Ca^2+^ burst can be explained by a combination of contributions from multiple biomolecules. As the reverse phenomenon of Ca^2+^ bursts during heating, the rate of Ca^2+^ uptake by the Ca^2+^-ATPase, SERCA, is reduced as the temperature is decreased, and the open probabilities of IP_3_R returns, i.e., is probably increased to the original level before heating. Then, the net Ca^2+^ flow (the amount of released Ca^2+^ per unit time) from the ER to the cytoplasm could be increased during recooling, and the Ca^2+^ burst is induced due to CICR in IP_3_R. However, when the cooling rate is relatively slow, the recovery rate of the net Ca^2+^ flow is also slow, which can be well compensated by the uptake activity of SERCA, exclusion pathways by other Ca^2+^ exchangers, and/or buffer function of Ca^2+^-binding proteins.

### Conclusions

This study was conducted to expand the abilities of the spatiotemporal manipulation of the temperature gradient for investigating cellular thermosensitivities and manipulating their functions. The results demonstrated that CGH images displayed on LCOS-SLM modulate the spatial phase pattern of IR light and generate predesigned temperature gradients at the microscopic focal plane. The holographic heating microscope visualized how thermosensitive Ca^2+^ signaling is generated and propagated in MDCK cells, rat hippocampal neurons, and rat neonatal cardiomyocytes. Moreover, the optical control of the temporal temperature gradient with the phase modulation of the IR light revealed the cooling-rate dependency of Ca^2+^ signaling in HeLa cells.

Although genetically encoded light-sensitive proteins are widely used in optogenetics for the optical manipulation of cellular functions, endogenous thermosensitive systems could be a target of optothermal manipulations ^7^. The applications of the holographic heating microscope will not be limited to thermosensitive Ca^2+^ signaling. Spatial temperature gradient affects cell morphologies such as membrane extension of HeLa cells ^23^ and neurite outgrowth ^32^ and cell division in *Caenorhabditis elegans* embryos ^11^. Therefore, the holographic heating microscope might be useful for manipulating cellular morphology, neural networks, and multicellular structures such as spheroids and organoids.

## Materials and Methods

### Optical setup of the holographic heating microscope

A polarized IR laser light (λ = 1,475 nm; AF4B150FK75L; Anritsu) was expanded and collimated with a collimator (F810FC-1550; Thorlabs). A reflective LCOS-SLM (SLM-100; Santec) with 1450×1050 pixels modulated the phase of light at each pixel. The irradiation was controlled with a mechanical shutter (SSH-25RA; Sigmakoki) and a controller (SSH-C2B; Sigmakoki). Linear polarizers (LPNIR050-MP2 and LPIREA100-C; Thorlabs) and plano-convex lenses (LA1951-C-ML, LA1608-C-ML, LA1131-C-ML, and LA1509-C-ML; Thorlabs) were used. An aluminum block was placed on a coverslip (S1112; Matsunami Glass). An inverted microscope (IX-83; Olympus) equipped with a four-wavelength high-power LED source (LED4D237; 365/490/565/660 nm; Thorlabs), a custom-made dichroic mirror for the IR laser light (Sigmakoki), and an EMCCD camera (iXon Life 888; Andor Technology) were used. CGH images were generated by the CGH generator (version 1.00.00, Santec).

### IR imaging and temperature measurement

Upconversion nanoparticles were synthesized by a thermal decomposition method using Na(CF_3_COO) and Ln(CF_3_COO)_3_ as precursors (Ln = Er^3+^ and Yb^3+^) ^21^. The emission spectrum excited by a 1,550 nm CW laser light (70.1 mW; 1550L-11A-NI-NT-NF; Integrated Optics) was measured with a spectrometer (SILVER-Nova 25 TEC ZAP-D; StellarNet). For IR imaging at the microscopic focal plane, the upconversion nanoparticles in chloroform were spread over a glass-bottom dish (3911–035; AGC Techno Glass) and dried at room temperature.

The temperature gradient was visualized with luminescent thermometer nanosheets, which were fabricated on a glass-bottom dish or a polymer-bottom dish (SF-T-D12; Fine Plus International) according to a previously reported method ^41^. The thermometer nanosheets were observed with an excitation filter (BP360-370, Olympus), a dichroic mirror (FF493/574-Di01; Semrock), an emission filter (FF01-512/630; Semrock), and a 10× (LMPLN10XIR; Olympus) or a 60× objective lens (PLAPON60XOTIRF; Olympus). The laser power of the IR light illuminated by the objective lenses was measured using a thermal power sensor at the sample stage (S175C; Thorlabs).

### Cell culture and Ca^2+^ imaging

Animal experiments were approved by the Institutional Animal Care and Use Committee of the National Institutes for Quantum Science and Technology. They were performed in accordance with the Fundamental Guidelines for Proper Conduct of Animal Experiments and Related Activities in Academic Research Institutions under the jurisdiction of the Ministry of Education, Culture, Sports, Science, and Technology of Japan.

MDCK cells (RCB0995) and HeLa cells (RCB0007) were purchased from the RIKEN BRC Cell Bank. Two days before the experiment, MDCK cells were transferred onto a polymer-bottom dish and cultured in minimum essential medium Eagle (MEM) (M5650; Sigma-Aldrich) containing 10% fetal bovine serum (FBS) (12483020; Thermo Fisher Scientific), 2 mM L-glutamine (G7513; Sigma-Aldrich), 100 U/mL penicillin, and 100 μg/mL streptomycin (15140122; Thermo Fisher Scientific) at 37°C in the presence of 5% CO_2_. HeLa cells on a glass-bottom dish were cultured in MEM containing 10% FBS (SH30910.03; HyClone), 2 mM L-glutamine, 100 U/mL penicillin, and 100 μg/mL streptomycin (10378016; Thermo Fisher Scientific).

Rat hippocampal neurons were obtained from 19-day embryos of Wistar rats (CLEA Japan) following our published protocol ^32^. Moreover, 4 ×10^5^ cells were cultured in 2 mL of culture medium composed of Neurobasal Plus Medium (A3582901; Thermo Fisher Scientific), B-27 Plus Supplement (A3582801; Thermo Fisher Scientific), 0.5 mM GlutaMAX Supplement (35050061; Thermo Fisher Scientific), 100 U/mL penicillin, and 100 μg/mL streptomycin on a polymer-bottom dish coated with 0.3 mg/mL collagen (Cellmatrix Type I-C; Nitta Gelatin) and 0.1 mg/mL poly-D-lysine (A-003-E; Merck) for 21 days at 37°C in the presence of 5% CO_2_. On day 2 after seeding, 1 μM cytosine-1-β-D(+)-arabinofuranoside (030-11951; FUJIFILM Wako Pure Chemical) was added. Half the volume of the culture medium was replaced twice in a week.

Cardiomyocytes were obtained from neonatal Wistar rats (Japan SLC) according to our published protocol (Shintani JGP 2014). Furthermore, 1.5 ×10^4^ cell suspension in 0.15 mL of the culture medium composed of Dulbecco’s MEM (08488-55; Nacalai Tesque), 10% FBS (Thermo Fisher Scientific), 4 mM L-glutamine (Sigma-Aldrich), 100 µM sodium pyruvate (P8574, Sigma-Aldrich), 100 U/mL penicillin, and 100 μg/mL streptomycin in a polymer-bottom dish that had been precoated with 0.3 mg/mL collagen was incubated overnight at 37°C in the presence of 5% CO_2_. From the next day, cells were cultured in 2 mL of the culture medium containing 100 μM 5-Bromo-2′-deoxyuridine (B5002; Sigma-Aldrich) for 3 days before the experiments.

On the day of the experiments, the cells were incubated in the culture medium containing 5 µM Cal-520, AM (21130; AAT Bioquest) for 90 min at 37°C in the presence of 5% CO_2_. After two washes with the imaging solution, cells in the imaging solution (140 mM NaCl, 5 mM KCl, 1 mM MgCl_2_, 1 mM Na_2_HPO_4_, 10 mM HEPES, 1.8 mM CaCl_2_, and 25 mM D(+)-glucose, pH 7.4, adjusted by NaOH) were set on the microscope stage for 10 min to stabilize the temperature before the observation. In the experiments for MDCK, neuron and HeLa cells, the temperature was set at 37°C using a thermostatically controlled incubator (INUC-KRi-F1; Tokai Hit). MDCK cells, neurons, and cardiomyocytes were observed with the 10× objective lens. HeLa cells were observed with the 60× objective lens. Cal-520 was excited by 490-nm light and observed with a mirror unit (U-FBNA; Olympus). Electrical stimulation (∼20 V/cm, 10 ms) was applied using an electronic stimulator (SEN-7203; Nihon Kohden) and an isolator (SS-104J; Nihon Kohden).

### Statistics

The Tukey–Kramer test was performed using OriginPro2021 (OriginLab). Data are expressed as mean ± SD.

## Author Contributions

K.O. and M.S. designed the research study; K.O., A.I., S.M., and T.G.O. performed the experiments; K.O. and M.S. analyzed the data. All authors interpreted the results and wrote the manuscript.

## Data Availability

The data supporting this article have been included as part of the Supplementary Information.

## Acknowledgments

We thank Ms. Noriko Uchida (National Institutes for Quantum Science and Technology) and Ms. Noriko Tawara (National Institutes for Quantum Science and Technology) for their technical assistance. This work was supported by the Japan Science and Technology Agency JPMJPR17P3 (to K.O.) and JPMJPR17P2 (to A.I.), JSPS KAKENHI Grant Numbers 22H05053 (to M.S.) and 22H05054 (to K.O.), Tokuyama Science Foundation (to A.I.), and Takeda Science Foundation (to M.S.).

