## Supplementary Information for "Spatiotemporal temperature control by holographic heating microscopy unveils cellular thermosensitive calcium signaling"

**Supplementary figure**

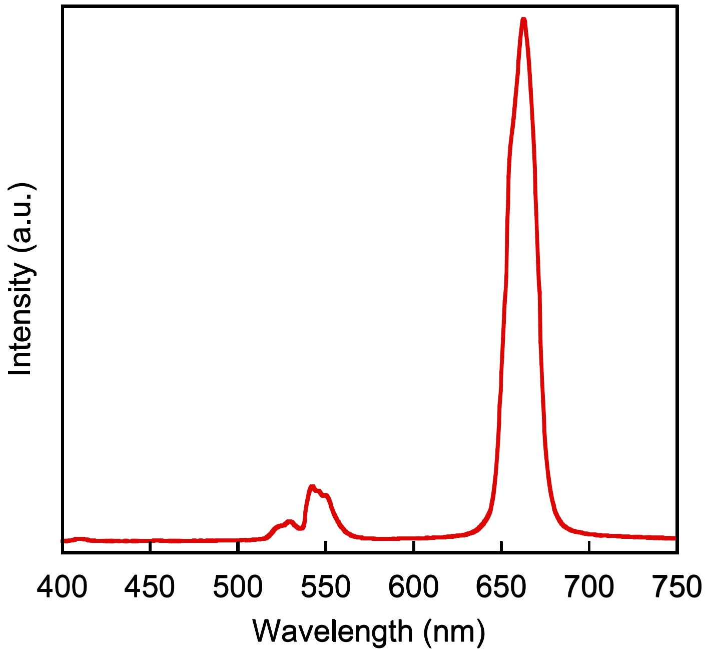


**Figure S1. Photoluminescence spectrum of the upconversion nanoparticles excited by 1,550-nm laser light.** The wavelength of the emission peak was 662 nm, originating from the upconversion emission of $Er^{3+}$.

**Supplementary movies**

**Movie S1. Spatiotemporal control of infrared laser light by the holographic heating microscope.** Heat pulses for 2 s were applied at different positions. Infrared (IR) laser light was visualized by an IR-viewer sheet with upconversion nanoparticles on a coverslip. Scale bar, 200 μm.

**Movie S2. Spatiotemporal control of the microscopic temperature by a holographic heating microscope.** Heat pulses for 2 s were applied at different positions. The temperature changes were visualized by a luminescence thermometer nanosheet on a polymer-bottom dish. Scale bar, 200 μm.

**Movie S3. Spatiotemporal manipulation of $Ca^{2+}$ signaling in MDCK by the holographic heating microscope.** Fluorescence images of Cal-520 in MDCK cells. Heat pulses for 10 s were initiated at 10, 30, and 50 s, respectively. The yellow arrows indicate the positions where the IR light was focused. Scale bar, 200 μm.

**Movie S4. Spatiotemporal manipulation of $Ca^{2+}$ signaling in rat hippocampal neurons using a holographic heating microscope.** Fluorescence images of Cal-520 in rat hippocampal neurons. Electrical stimulation (ES) was applied at 10 s. Heat pulses for 2 s were initiated at 20, 30, and 40 s. The yellow arrows indicate the positions where the IR light was focused. Scale bar, 200 μm.

**Movie S5. Holographic heating microscope accelerated the beating of neonatal rat cardiomyocytes.** Fluorescence images of Cal-520 in rat neonatal cardiomyocytes. Heat pulses for 20 s were initiated at 20 and 60 s. The yellow arrows indicate the positions where the IR light was focused. Scale bar, 200 µm.

**Movie S6. $Ca^{2+}$ dynamics in HeLa cells responding to rapid recooling.** Fluorescence images of Cal-520 in HeLa cells. Optical heating for 30 s was initiated at 10 s. The yellow arrows indicate the positions where the IR light was focused. Rapid recooling for 3 s was initiated at 40 s. Scale bar, 50 µm.

**Movie S7. $Ca^{2+}$ dynamics in HeLa cells responding to middle recooling.** Fluorescence images of Cal-520 in HeLa cells. Optical heating for 30 s was initiated at 10 s. The yellow arrows indicate the positions where the IR light was focused. Middle-rate recooling for 7 s was initiated at 40 s. Scale bar, 50 µm.

**Movie S8. $Ca^{2+}$ dynamics in HeLa cells in response to slow cooling.** Fluorescence images of Cal-520 in HeLa cells. Optical heating for 30 s was initiated at 10 s. The yellow arrows indicate the positions where the IR light was focused. Slow recooling for 14 s was initiated at 40 s. Scale bar, 50 µm.
